# Dynamic 11C-PiB PET shows cerebrospinal fluid flow alterations in Alzheimer’s disease and multiple sclerosis

**DOI:** 10.1101/493734

**Authors:** Julia J. Schubert, Mattia Veronese, Livia Marchitelli, Benedetta Bodini, Matteo Tonietto, Bruno Stankoff, David J. Brooks, Alessandra Bertoldo, Paul Edison, Federico E. Turkheimer

**Affiliations:** Department of Neuroimaging, IoPPN, King’s College London, London, United Kingdom; Sorbonne Universités, UPMC Paris 06, Institut du Cerveau et de la Moelle épinière, ICM, Hôpital de la Pitié Salpêtrière, Paris, France; Imperial College London, London, United Kingdom; Department of Information Engineering, University of Padova, Padova, Italy

**Keywords:** cerebrospinal fluid, glymphatic system, PiB PET, Alzheimer’s disease, multiple sclerosis

## Abstract

Cerebrospinal fluid (CSF) plays an important role in the clearance of solutes and maintenance of brain homeostasis. ^11^C-PiB PET was recently proposed as a tool for detection of CSF clearance alterations in Alzheimer’s disease. The current study seeks to investigate the magnitude of ^11^C-PiB PET signal in the lateral ventricles of an independent group of Alzheimer’s and mild cognitive impairment subjects. We have also evaluated multiple sclerosis as a model of disease with CSF clearance alterations without amyloid-beta tissue accumulation.

**Methods:** A set of Alzheimer’s and mild cognitive impairment subjects and a set of multiple sclerosis subjects with matched healthy controls underwent MRI and dynamic ^11^C-PiB PET. Manual lateral ventricle regions of interest were generated from MRI data. PET data was analysed using a simplified reference tissue model with cerebellum or a supervised reference region, for the Alzheimer’s and multiple sclerosis datasets, respectively. Magnitude of ^11^C-PiB signal in the lateral ventricles was calculated as area under curve from 35 to 80 minutes and standard uptake value ratio (SUVR) from 50 to 70 minutes. Compartmental modelling analysis was performed on a separate dataset containing Alzheimer’s and matched healthy control data with an arterial input function to further understand the kinetics of the lateral ventricular ^11^C-PiB signal.

**Results:** Analysis of variance revealed significant group differences in lateral ventricular SUVR across the Alzheimer’s, mild cognitive impairment, and healthy control groups (p=0.004). Additional pairwise comparisons revealed significantly lower lateral ventricular SUVR in Alzheimer’s compared to healthy controls (p<0.001) and mild cognitive impairment (p=0.029). Lateral ventricular SUVR was also significantly lower in multiple sclerosis compared to healthy controls (p=0.008). Compartmental modelling analysis revealed significantly lower uptake rates of ^11^C-PiB signal from blood (p=0.005) and brain tissue (p=0.004) to the lateral ventricles in Alzheimer’s compared to healthy controls. This analysis also revealed significantly lower clearance of ^11^C-PiB signal out of the lateral ventricles in Alzheimer’s compared to healthy controls (p=0.002).

**Conclusion:** Overall, these results indicate that dynamic ^11^C-PiB PET can be used to observe pathological changes in cerebrospinal fluid dynamics and that cerebrospinal fluid-mediated clearance is reduced in Alzheimer’s disease and multiple sclerosis compared to healthy controls.

## Introduction

There has been great interest surrounding cerebrospinal fluid (CSF) dynamics since the existence of the glymphatic system was proposed in 2012 (*1–4*). The glymphatic system has been suggested as being largely responsible for the clearance of waste from the brain (*1,5*). The original description of the glymphatic system proposed that CSF penetrates the brain via para-arterial spaces, enters the brain parenchyma where it combines with interstitial fluid (ISF) and collects waste and other solutes, and returns to the subarachnoid space or clears through the vascular and lymphatic systems via paravenous spaces (Fig. 1A). This system is thought to be analogous to the body’s peripheral lymphatic system with the additional involvement of glial cells (*1*). Although there is still debate over the exact mechanism underlying the glymphatic system (*2,3,6–8*), there is an overall agreement that a para- and/or peri-vascular clearance system of the brain exists and that it is closely linked to the production and flow of CSF.

**Figure 1.**
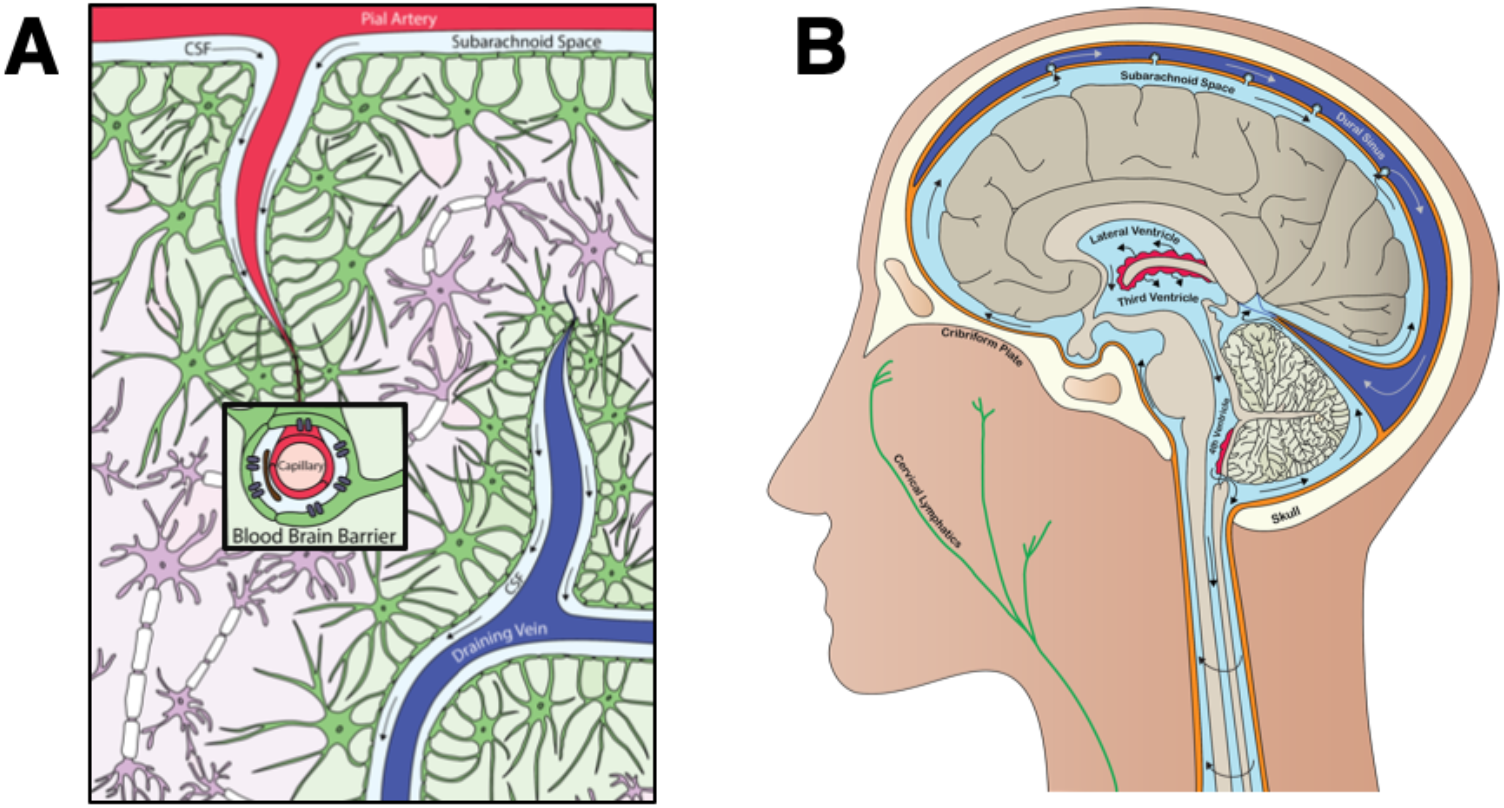
Interrelationship between clearance and fluid systems of the brain. **A**. The glymphatic system. **B**. The cerebrospinal fluid system. Illustrations by Julia J Schubert.

CSF is predominantly produced by the choroid plexuses of the lateral, third, and fourth ventricles. Interstitial fluid of surrounding brain tissues, excluding those of the circumventricular organs, is able to exchange quite freely with ventricular CSF due to the presence of gap junctions in the ependymal cell lining of the adult ventricular system that allow free diffusion of small molecules (*9,10*). CSF generally has a net positive flow through the ventricular system and out to the subarachnoid space that surrounds the brain and spinal cord (Fig. 1B). CSF is eventually cleared to the venous system through arachnoid villi, to the cervical lymphatics through the cribriform plate (*11*), and to the recently discovered meningeal lymphatics that line the dural sinuses (*12*). The trans-astrocytic water movements associated with CSF flow into and CSF/ISF flow out of the brain parenchyma are facilitated by aquaporin-4 (AQP4) water channels, which are localized along perivascular astrocytic end-feet that contribute to the blood brain barrier (*1*). Alterations to the normal flow of CSF in the brain could be a contributing factor to the accumulation of toxic substances in the brain interstitium and may be related to the pathogenesis of neurological disorders such as Alzheimer’s disease and multiple sclerosis.

The glymphatic system has been shown to be predominantly responsible for the clearance of amyloid-beta (A*β*) (*1,13*), the main component of the amyloid plaques found in Alzheimer’s disease brain. Significant accumulation of A*β* has been observed after inhibition of glymphatic transport (*13*) and deletion of the AQP4 gene suppresses clearance of soluble A*β* (*1*). These findings suggest that inhibition of CSF flow could contribute to glymphatic dysfunction and the development of extracellular A*β* plaques characteristic of Alzheimer’s disease. Phase-contrast MRI has also shown significantly decreased CSF flow in multiple sclerosis compared to healthy controls (*14,15*) and associations have been observed between decreased CSF flow and conversion rate from clinically isolated syndrome to clinically defined multiple sclerosis (p=0.007) and relapse rate in relapsing-remitting multiple sclerosis (p=0.035) (*14*). Further, loss of perivascular AQP4 localization has been observed in an experimental autoimmune encephalomyelitis mouse model of multiple sclerosis (*16*). These previous findings further support a link between CSF flow alterations and neurological diseases such as Alzheimer’s disease and multiple sclerosis.

Despite the current interest in CSF dynamics, non-invasive in-vivo methods for measuring glymphatic function are limited and the majority of existing data has been collected using animal models (*5,13,16*). ^11^C-PiB is a neutral lipophilic benzothiazole PET tracer and is used to image A*β* plaques in Alzheimer’s disease (*17*) and, more recently, to quantify in vivo myelin loss and regeneration in multiple sclerosis (*18*). ^11^C-PiB PET has also recently been used to quantify CSF clearance in humans (*19*). Small molecule radiotracers enter the CSF of the lateral ventricles either directly from the blood through a thick layer of epithelial cells that make up the choroid plexuses or from the brain parenchyma via diffusion through the ependymal cells that line the adult ventricular system (*10*). Given the close connection between CSF flow and the glymphatic system, ^11^C-PiB PET could also serve as an in-vivo marker of glymphatic system function. Here we aim to replicate previous results that showed decreased lateral ventricular ^11^C-PiB signal magnitude in Alzheimer’s disease compared to controls (*19*) and to extend the method to mild cognitive impairment and multiple sclerosis patient groups. Based on previous research, each of these patient groups are expected to have altered CSF dynamics. However, due to differences in the pathogenesis of Alzheimer’s disease compared to multiple sclerosis, A*β* deposition is only expected in the Alzheimer’s disease and mild cognitive impairment groups. Inclusion of the multiple sclerosis group allows for further testing of our dynamic PET method for measuring CSF dynamics without the confounding factor of A*β* accumulation in brain tissue. We also aim to further understand the kinetics of the ^11^C-PiB PET signal in the lateral ventricles by performing compartmental modelling analysis. Combined, these results will allow us to make appropriate interpretations about the lateral ventricular ^11^C-PiB signal magnitude and how these measurements may relate to CSF clearance and glymphatic function in healthy and diseased populations.

## Materials and Methods

### Participants

To investigate differences in CSF clearance in Alzheimer’s disease and multiple sclerosis, analysis was performed on two datasets. One dataset included 11 Alzheimer’s disease patients (6/5 women/men; mean age: 66.6 ± 4.4 years), 12 mild cognitive impairment patients (4/8 women/men; mean age: 68.6 ± 7.9 years) and 12 age- and sex-matched healthy controls (5/7 women/men; mean age: 64.4 ± 6.6 years) that were recruited from Hammersmith Hospital NHS Trust, the National Hospital for Neurology and Neurosurgery, St. Margaret’s Hospital, and Victoria Hospital, UK. Recruitment of the healthy controls in this dataset was aided by enrolment of spouses of the Alzheimer’s disease subjects. The Alzheimer’s disease and healthy control data has previously been reported alone (*20*) and with comparison to the mild cognitive impairment data (*21*). Clinically probable diagnoses of Alzheimer’s disease were assigned based on the National Institute of Neurological and Communicative Diseases and Stroke/AD and Related Disorders Association (NINCDS-ADRDA) and Diagnostic and Statistical Manual of Mental Disorders (DSM-IV) criteria. All Alzheimer’s subjects also met the National Institute on Aging/Alzheimer’s Association (NIA/AA) criteria. Alzheimer’s subjects aged 55 to 79 with a clinical diagnosis of Alzheimer’s disease before enrolment were included in the study. Patients and controls with history of mental health issues, significant white matter microvascular disease on MRI, a contraindicative MRI, a history of drug or alcohol abuse, and/or any other neurological causes were excluded. For full details on inclusion and exclusion criteria for the Alzheimer’s subjects, please refer to the original report of this dataset (*20*). Mild cognitive impairment subjects were chosen based on similar exclusion criteria as the Alzheimer’s subjects and all mild cognitive impairment subjects fulfilled Petersen’s criteria for amnestic mild cognitive impairment.

A second dataset included 20 relapsing-remitting multiple sclerosis patients (13 women; mean age: 32.3 ± 5.6 years) and eight age- and sex-matched healthy controls (5 women; mean age: 31.6 ± 6.3 years). All patients were diagnosed with multiple sclerosis according to the revised McDonald criteria (*22*) and had at least one gadolinium-enhancing lesion when they entered the study. Additional information on inclusion and exclusion criteria can be found in the original report of this dataset (*18*). The imaging protocols were approved by the local ethics committees and all participants gave written informed consent prior to data collection.

### Clinical Assessments

#### Alzheimer’s disease/mild cognitive impairment (AD/MCI) dataset

Detailed neurological assessments were performed for nine Alzheimer’s subjects. These assessments included taking a patient history from a close relative, routine blood analysis, and EEG. The following neuropsychometric assessments were also performed: Mini-Mental State Examination (MMSE) (*23*), Warrington short recognition memory tests (WRTM) for words and faces, Alzheimer’s Disease Assessment Scale Word List Learning test and 30 minute delayed recall (*24*), immediate and delayed recall of modified complex figure (*25*), Digit Span forwards (*26*), Trail Making Part A (*27*), clock drawing (*28*), copy of modified complex figure (*25*), 30-item Boston Naming Test (*29*), letter fluency (FAS) (*30*), and category fluency (animals, birds, and dogs). All mild cognitive impairment subjects also underwent a comprehensive assessment that included neurologic examination, neuropsychological testing, and MRI.

#### Multiple sclerosis dataset

Measures of disease severity including Expanded Disability Status Scale (EDSS) (*31*) and the Multiple Sclerosis Severity Scale (MSSS) (*32*) were used to rate nineteen multiple sclerosis subjects at the time of enrolment.

### PET and MRI data acquisition

#### AD/MCI dataset

All subjects had 90-minute ^11^C-PiB PET on a Siemens ECAT EXACT HR+ scanner with 3D acquisition and an axial field of view of 15.5 cm. All subjects were given an intravenous bolus injection of ^11^C-PiB (mean=365 ± 24 MBq) at the start of each scan. Image reconstruction and data processing, including scatter correction, was performed using standard Siemens software. All subjects also had MRI, which was performed with a 1.5 Tesla GE scanner. The T1-weighted structural MRI data was used for ROI segmentation in this work. Additional information on the PET and MRI protocols can be found in the original reports of this data (*20,21*).

#### Multiple sclerosis dataset

A high-resolution tomograph (HRRT; CPS Innovations, Knoxville, TN) was used to perform 90-minute ^11^C-PiB PET on all subjects. All subjects were given an intravenous bolus injection of ^11^C-PiB (mean=358 ± 34 MBq) at the start of each scan. An intraslice spatial resolution of about 2.5 mm full width at half maximum was achieved, with 25-cm axial and 31.2-cm transaxial fields of view. Poisson ordered subset expectation maximization algorithm with 10 iterations was used for image reconstruction. To assist in reducing the effects of partial volume in the multiple sclerosis dataset (*33*), the reconstructed images were smoothed with a filter implementing point spread function. All subjects also underwent MRI scanning on a 3 Tesla Siemens TRIO 32-channel TIM system. T1-weighted structural MRI data collected before gadolinium injection was used for ROI segmentation in this work. The PET image acquisition, reconstruction, and quantification techniques have been previously described (*34*). Additional information on the PET and MRI protocols can be found in the original report of this dataset (*18*).

### Data analysis

#### AD/MCI dataset

Preprocessing of the PET and MRI data from the AD/MCI dataset was performed and time activity curves (TACs) were generated using MIAKAT™ (version 4.2.6) (*35*) software. MIAKAT™ is implemented in MATLAB (version R2015b; The MathWorks, Inc., Natick, Mass.) and the preprocessing pipeline uses tools from SPM12 and FSL (version 5.0.9) (*36*) analysis toolboxes to perform brain extraction, tissue segmentation, rigid and nonlinear registration to an MNI template (*37*), region of interest (ROI) definition, and motion correction. An additional gray matter ROI was defined that excluded the cerebellum reference region. One Alzheimer’s and one mild cognitive impairment subject did not pass quality control due to misregistration artifacts and their data were later excluded from the AD/MCI analysis.

#### Multiple sclerosis dataset

Motion correction of the multiple sclerosis dataset was performed by realigning each PET frame to a common reference space, as has been previously described (*38*). T1-weighted MRI images were registered to the ^11^C-PiB PET. A priori designated ROIs were used for a supervised reference region and a gray matter ROI used in the multiple sclerosis analysis. The gray matter ROI excluded reference and cerebellar gray matter for comparison with the Alzheimer’s dataset. TAC extraction for the multiple sclerosis dataset was performed using in-house MATLAB™ (version R2017a; The MathWorks, Inc., Natick, Mass.) scripts.

#### Both datasets

Manual lateral ventricle ROIs were generated for all subjects using the subject T1-weighted structural MRI data and the ITK-SNAP (*39*) (itksnap.org) snake tool, following previously described guidelines for lateral ventricle extraction (*40*). The lateral ventricle ROIs were then eroded by two voxels (5.2 mm) using the erode function given by FSL’s ‘fslmaths’ utility package (version 5.0.9) (*36*) in order to reduce partial volume effects of the surrounding tissues. An example lateral ventricle ROI is shown in Figure 2.

**Figure 2.**
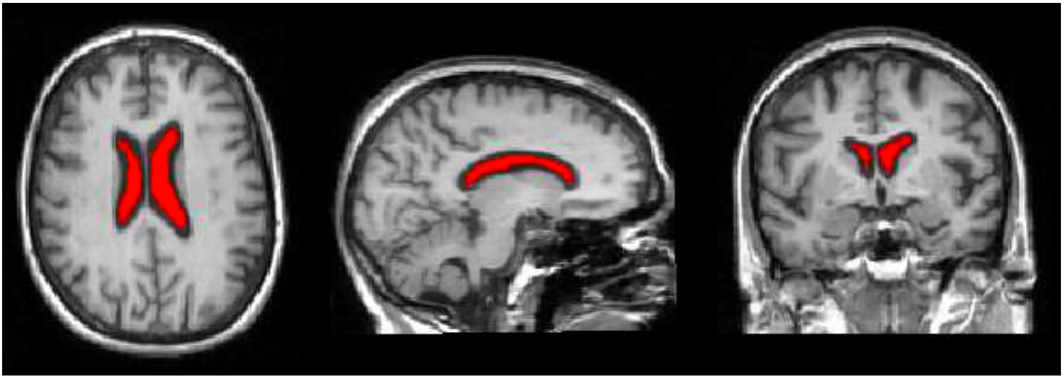
Lateral ventricle region of interest (ROI). Lateral ventricle cerebrospinal fluid is filled with red. ROI was generated using ITK-SNAF snake tool and has subsequently been eroded by 2 voxels (5.2mm).

Standardized uptake value ratios (SUVRs) from 50 to 70 minutes were calculated for the gray matter and lateral ventricle ROIs for each subject from both datasets using cerebellum or a supervised reference region for the AD/MCI and multiple sclerosis datasets, respectively. For consistency with a previous study (*19*), area-under-curve (AUC) from 35 to 80 minutes was also calculated for the lateral ventricle ROI for a subset of the AD/MCI dataset. AUC_35-80_ was also calculated for the lateral ventricle ROI for all subjects in the multiple sclerosis dataset.

### Compartmental modelling analysis

#### Participants

A dataset that included 11 Alzheimer’s disease subjects and 11 age- and sex-matched healthy controls was included for sole use in compartmental modelling analysis. This data has been previously reported (*41*). All Alzheimer’s subjects in this dataset met the NINCDS-ADRDA criteria for clinical probable Alzheimer’s disease and had MMSE scores greater than 16 at study inclusion. Healthy controls were recruited by advertisement or during participation in a separate long-term follow-up study in the clinic. The imaging protocol was approved by the local ethics committee and all participants gave written informed consent prior to data collection.

#### PET Data Acquisition

All subjects underwent 90-minute ^11^C-PiB PET with online arterial sampling. PET data was collected on an ECAT EXACT H+ scanner (Siemens/CTI). Metabolite-corrected arterial input functions were determined using the measured percentage of radioactive parent compound in plasma. Further details about the PET protocol can be found in the original report of this dataset (*41*).

#### Compartmental model

The final model used to describe ^11^C-PiB PET kinetics in the lateral ventricles is shown in Fig. 3. This model includes two compartments to account for the signal from both bound and unbound ventricle pools and two input functions. One input function corresponds to the whole brain gray matter, which describes the tracer transport from the brain tissue into the ventricles, and one input function corresponds to the parent plasma arterial input function, which describes the tracer transport from the blood into the lateral ventricles. This model was determined based on known biological restraints of the CSF system and is also consistent with a previously presented compartmental model of the CSF clearance system (*42*).

**Figure 3.**
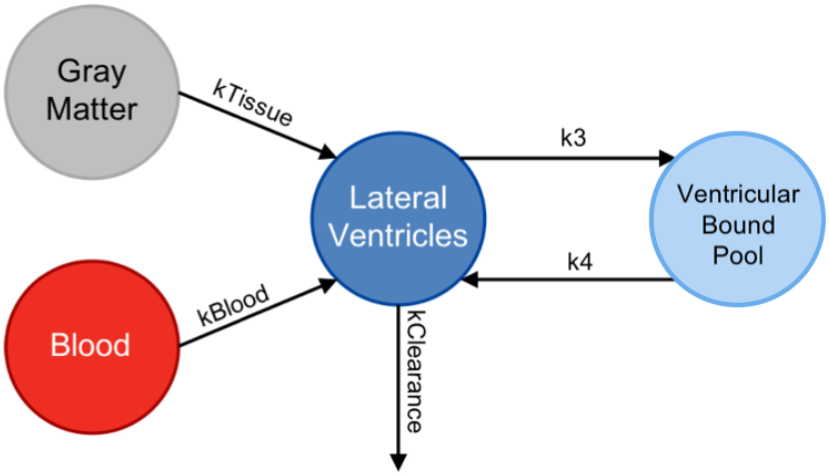
Kinetic model used in compartmental modelling analysis. *Kclearance* includes total clearance from lateral ventricles to blood, tissue, and rest of ventricular system.

#### Data Analysis

Compartmental modelling analysis was performed with SAAM II software (*43*). The lateral ventricle ROI used for TAC extraction was defined using a registered MNI atlas and the ITK-SNAP (*39*) (www.itksnap.org) snake tool with additional manual drawing when necessary. This particular dataset was used because it included an arterial input function necessary for accurate signal measurements from the blood, which is required for compartmental modelling analysis. The rate constants *Kclearance, K1tissue, K1blood, k3*, and *k4* represent the rate of ^11^C-PiB signal between tissues in the system. *Kclearance* takes into account the clearance of signal from the lateral ventricles to blood, surrounding tissues, and the rest of the ventricular system because it is not possible to differentiate between these clearance pathways. The SAAM II software was used to iteratively fit the TAC data to the model and final values of the rate constants were recorded for each subject.

### Statistical analyses

All statistical analyses were performed in SPSS (version 24.0, Chicago, IL). Shapiro-Wilk’s W test was used to test for normality of the data. Analysis of variance was used to investigate group differences in lateral ventricle and gray matter ^11^C-PiB signal in the AD/MCI dataset. Paired differences in lateral ventricle ^11^C-PiB SUVRs between groups in the AD/MCI and multiple sclerosis datasets were investigated using independent-samples t-tests. Paired differences in lateral ventricle ^11^C-PiB AUC_35-80_ between groups were investigated using independent samples t-tests for the AD/MCI dataset and using the Mann-Whitney U test for the multiple sclerosis dataset. Paired differences in gray matter ^11^C-PiB SUVRs between groups were investigated using the Mann-Whitney U test for the AD/MCI dataset and using an independent samples t-test for the multiple sclerosis dataset. Spearman’s correlation was used to investigate the relationship between gray matter and lateral ventricular ^11^C-PiB SUVRs as well as between clinical scores and lateral ventricular ^11^C-PiB signal in Alzheimer’s disease and multiple sclerosis patient groups. Group differences in *Kclearance, k3*, and *k4* were investigated using independent-samples t-tests. Group differences in *K1tissue* and *K1blood* were investigated using the Mann-Whitney U test. Brain size and ventricle size were tested as possible covariates.

## Results

### Lateral ventricular ^11^C-PiB signal magnitude in relation to diagnosis

Analysis of variance (ANOVA) revealed significant group differences in lateral ventricle SUVRs across the Alzheimer’s disease, mild cognitive impairment, and healthy control groups (F(2, 30)=6.86, p=0.004), which remained significant when corrected for ventricle size (F(2, 29)=3.34, p=0.050). Brain size was not found to be a significant covariate in the AD/MCI dataset. Additional pairwise comparisons revealed significantly lower magnitude of lateral ventricular ^11^C-PiB, as measured by SUVR, in Alzheimer’s disease (M=0.37, SD=0.09) compared to healthy controls (M=0.57, SD=0.15) (t(*18*)=4.08, p<0.001) and in Alzheimer’s disease compared to mild cognitive impairment (M=0.49, SD=0.14) (t(*19*)=2.36, p=0.029). No significant difference in lateral ventricle SUVR was observed between mild cognitive impairment and healthy controls (t(*21*)=1.37, p=0.185) (Fig. 4A). Lateral ventricle AUC_35-80_ was significantly lower in Alzheimer’s disease (M=0.47, SD=0.13) than in healthy controls (M=0.72, SD=0.20) (t(*18*)=3.32, p=0.004) and mild cognitive impairment (M=0.65, SD=0.18) (t(*17*)=2.47, p=0.024). ANOVA also revealed significant group differences in lateral ventricle AUC_35-80_ across the Alzheimer’s disease, mild cognitive impairment, and healthy control groups (F(2,26)=5.53, p=0.010) (Fig. 4B).

**Figure 4.**
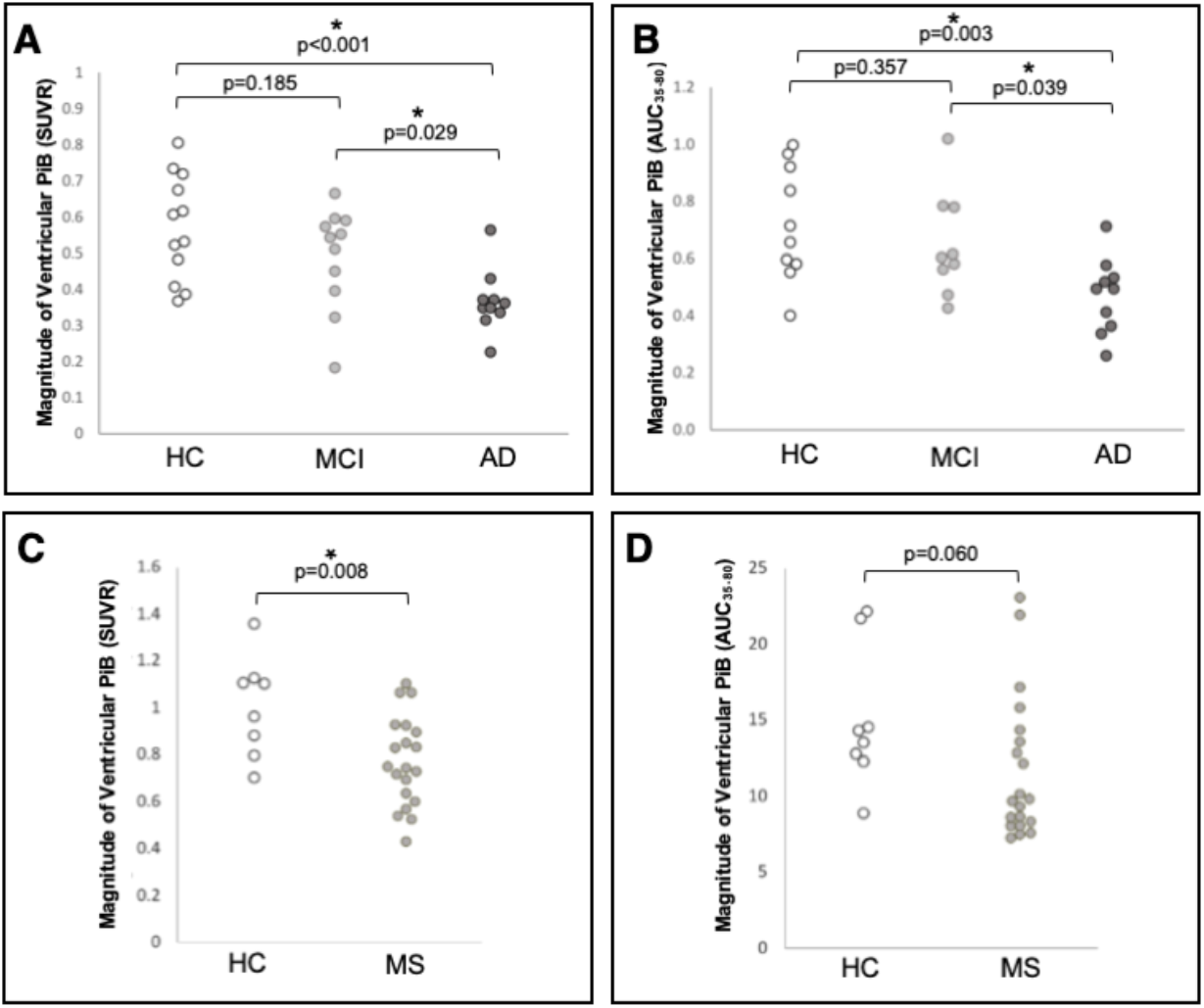
Group differences in lateral ventricle ^11^C-PiB measurements. **A**: Lateral ventricle ^11^C-PiB standard uptake value ratios (SUVR) in healthy controls (HC), mild cognitive impairment (MCI), and Alzheimer’s disease (AD). **B**: Lateral ventricle ^11^C-PiB AUC_35-80_ in HC, MCI, and AD. **C**: Lateral ventricle ^11^C-PiB SUVR in HC and multiple sclerosis (MS). **D**: Lateral ventricle ^11^C-PiB AUC_35-80_ in HC and MS.

Magnitude of lateral ventricle ^11^C-PiB, as measured by SUVR, was significantly lower in multiple sclerosis (M=0.77, SD=0.19) than in healthy controls (M=1.01, SD=0.21) (t(*26*)=2.87, p=0.008), which remained significant when corrected for ventricle size (F(1, 25)=4.35, p=0.047) (Fig. 4C). Brain size was not found to be a significant covariate in the multiple sclerosis dataset. There were no other significant differences in AUC_35-80_ measurements between groups.

### Correlations with tissue ^11^C-PiB

We observed significant negative correlations in ^11^C-PiB signal, as measured by SUVR, between gray matter and lateral ventricle ROIs in mild cognitive impairment (r=−0.664, p=0.026) and AD/MCI-matched control (r=−0.909, p<0.001) groups, which was not observed in Alzheimer’s disease (r=−0.479, p=0.162) (Fig. 5).

**Figure 5.**
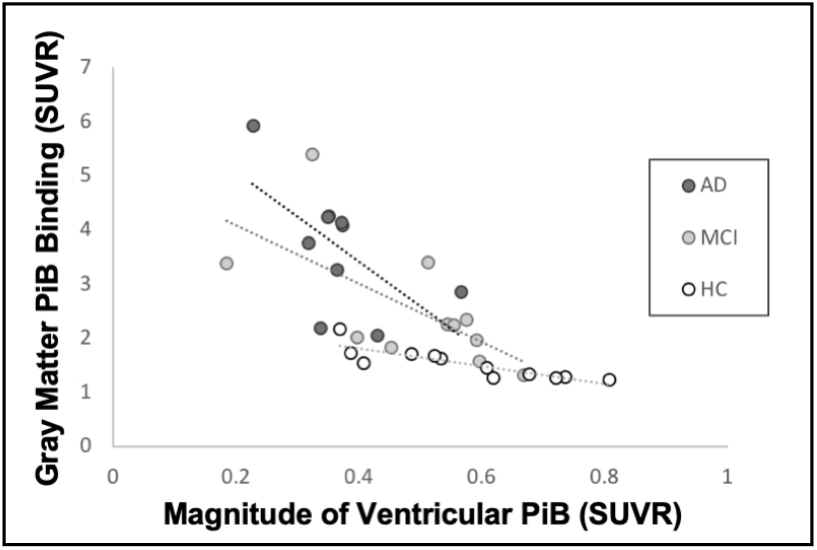
Gray matter and lateral ventricle ^11^C-PiB signal correlations. Correlations between gray matter ^11^C-PiB binding and magnitude of lateral ventricle ^11^C-PiB in Alzheimer’s disease (AD), mild cognitive impairment (MCI), and healthy controls (HC). All signal measurements are calculated as standard uptake value ratios (SUVR). AD: r=−0.479, p=0.162; MCI: r=−0.664, p=0.026; HC: r=−0.909, p<0.001

We did not observe any significant correlations in SUVRs between gray matter and lateral ventricle regions in the multiple sclerosis (r=0.045, p=0.850) or multiple sclerosis-matched control (r=0.333, p=0.420) groups.

### Gray matter ^11^C-PiB signal magnitude in relation to diagnosis

ANOVA revealed significant group differences in gray matter SUVRs across Alzheimer’s disease, mild cognitive impairment, and healthy control groups (F(2,30)=14.53, p<0.001). Gray matter ^11^C-PiB signal, as measured by SUVR, was observed to be significantly higher in Alzheimer’s disease (M=3.68, SD=1.14) compared to healthy controls (M=1.53, SD=0.28) (U=119.0, p<0.001) and mild cognitive impairment (M=2.52, SD=1.15) (U=87.0, p=0.024) and in mild cognitive impairment compared to healthy controls (U=116.0, p=0.002) (Fig. 6A). Gray matter ^11^C-PiB SUVRs were not significantly different between multiple sclerosis and healthy controls (t(*26*)=1.32, p=0.198) (Fig. 6B).

**Figure 6.**
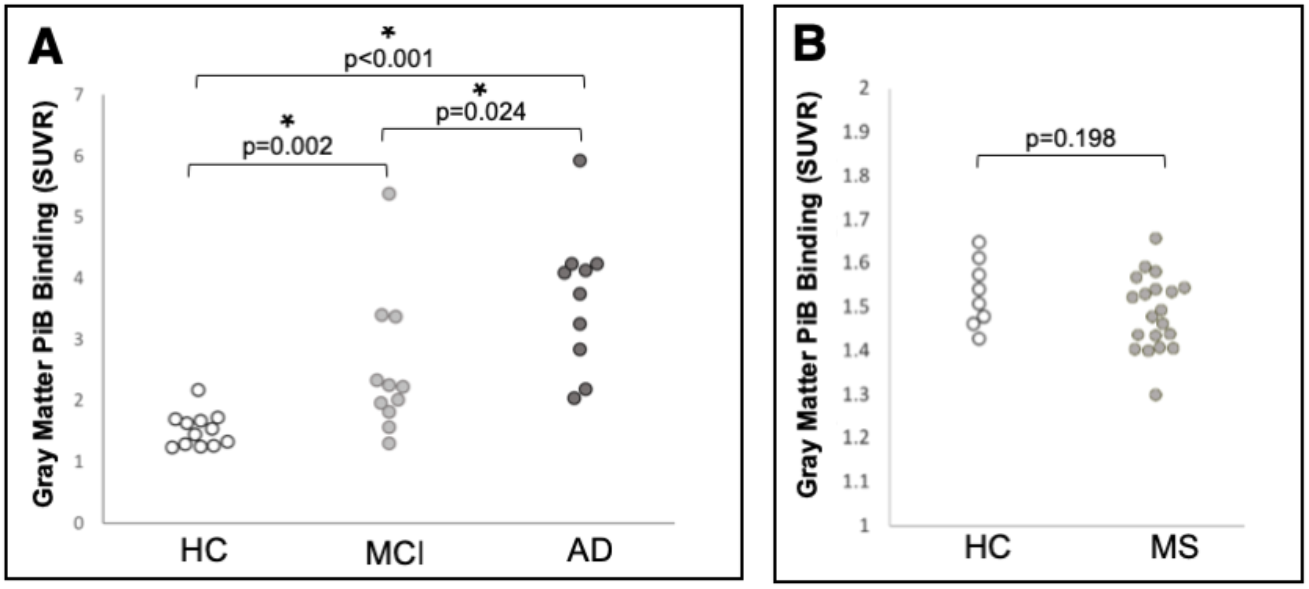
Group differences in gray matter ^11^C-PiB standard uptake value ratios (SUVR). **A**: Gray matter ^11^C-PiB SUVR in healthy controls (HC), mild cognitive impairment (MCI), and Alzheimer’s disease (AD). **B**: Gray matter ^11^C-PiB SUVR in HC and multiple sclerosis (MS).

### Correlations with disease severity measures

#### AD/MCI dataset

We did not observe any significant correlations between MMSE and lateral ventricle SUVRs in the Alzheimer’s disease group (r=0.518, p=0.153). No other significant correlations were observed between disease severity measures and lateral ventricle ^11^C-PiB measures in the Alzheimer’s disease group.

#### Multiple sclerosis dataset

We did not observe any significant correlations between MSSS (r=0.075, p=0.762), EDSS (r=−0.003, p=0.991), or disease duration (r=−0.144, p=0.555) and lateral ventricle SUVRs. We observed significant correlations between lesion load, as defined by the volume of white matter lesions, and lateral ventricle SUVRs (r=−0.493, p=0.032) and AUC_35-80_ (r=−0.595, p=0.007) (Fig. 7). We did not observe any other statistically significant correlations between disease severity measures and lateral ventricle ^11^C-PiB signal measures in the multiple sclerosis group.

**Figure 7.**
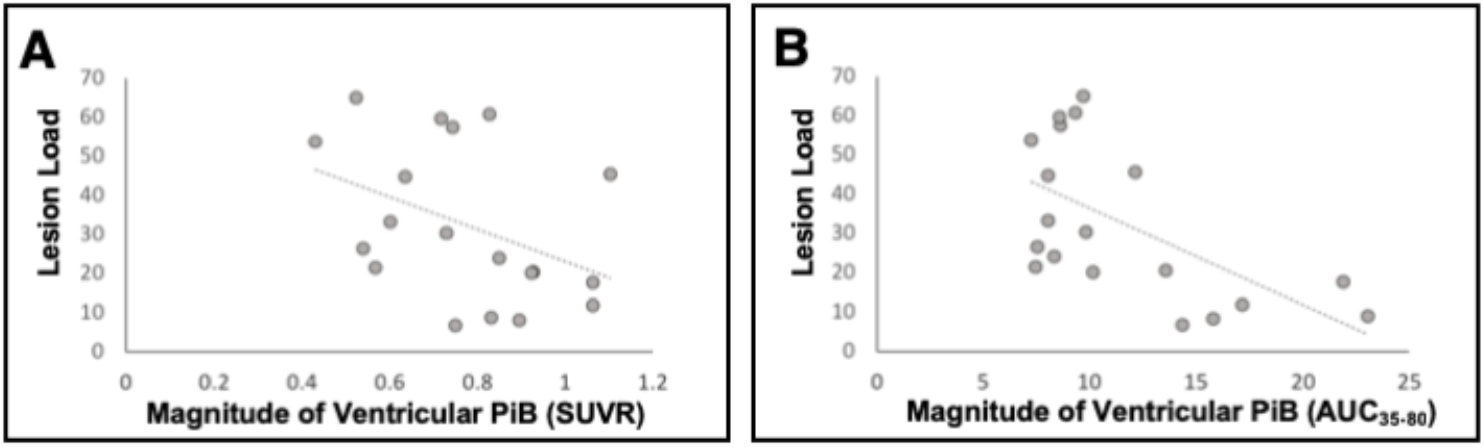
Lateral ventricle ^11^C-PiB measures and lesion load correlations. **A**: Correlation between lesion load and lateral ventricle standard uptake value ratios (SUVR) in multiple sclerosis (MS) (r=−0.493, p=0.032). **B**: Correlation between lesion load and lateral ventricle AUC_35-80_ in MS (r=−0.595, p=0.007). Lesion load is defined as volume of white matter lesions.

### Compartmental modelling analysis

The final compartmental model used in our analysis was able to reliably fit the lateral ventricle TAC data. Example fits of the lateral ventricle TAC data from one healthy control and one Alzheimer’s disease subject using the final compartmental model are shown in Fig. 8.

**Figure 8.**
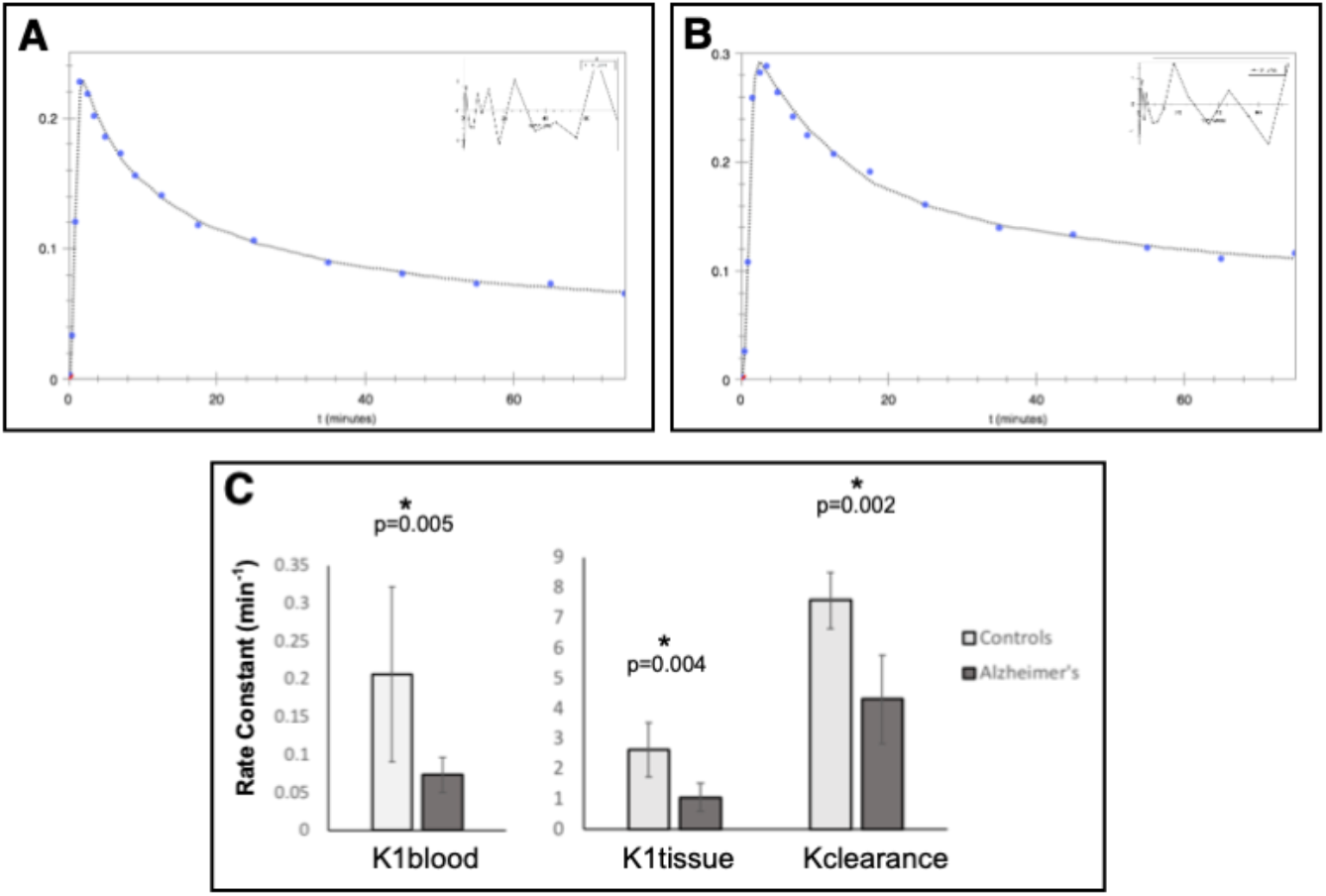
Example data fits and results from compartmental modelling. **A**: healthy control lateral ventricle TAC data fit example. **B**: Alzheimer’s patient lateral ventricle TAC data fit example. The weighted residuals are shown in the upper right corner of each image. **C**: Results from compartmental modelling analysis. Healthy controls and Alzheimer’s group differences in *Klblood, Kltissue*, and *Kclearance* rates. Error bars represent 95% confidence intervals.

Results from compartmental modelling analysis revealed significant group differences in rates of ^11^C-PiB signal exchange across tissues between Alzheimer’s disease and healthy control subjects. The rate of signal from blood to the lateral ventricles (*K1blood*) (U=13.0, p=0.005), from tissue to lateral ventricles (*K1tissue*) (U=12.0, p=0.004), and from lateral ventricles to blood, tissue, and the rest of the ventricular system (*Kclearance*) (t(*18*)=3.70, p=0.002) were significantly lower in Alzheimer’s disease compared to healthy controls (Fig. 8C). We did not observe any other significant differences between groups.

Simplified compartmental models, including one without specific binding in the lateral ventricles and one without a brain tissue input function, were also compared to the final compartmental model used in our analysis. The final model used was superior to simplified models, as determined by comparison of Akaike information criterion scores.

## Discussion

The observation of decreased ^11^C-PiB signal in the lateral ventricle ROIs in Alzheimer’s disease and multiple sclerosis patient groups compared to healthy controls indicates that dynamic PET measures can be used to observe pathological changes in CSF dynamics. We have successfully replicated a previous study that showed decreased lateral ventricle ^11^C-PiB signal in Alzheimer’s disease compared to healthy controls. Here, we have also shown that lateral ventricle ^11^C-PiB is significantly lower in multiple sclerosis compared to healthy controls and that mild cognitive impairment subjects have intermediate ^11^C-PiB measures between those of healthy controls and Alzheimer’s disease. We have also replicated previous results showing a significant inverse correlation between gray matter and lateral ventricle ^11^C-PiB binding in Alzheimer’s disease and our results show that this negative correlation is present in mild cognitive impairment as well as in AD/MCI-matched healthy controls. Our compartmental modelling analysis reveals that the reduced lateral ventricle signal is likely due to less tracer entering the lateral ventricles through the blood and through the brain tissue in Alzheimer’s disease compared to healthy controls. This analysis also showed that tracer is cleared from the lateral ventricles out to the surrounding tissues and the ventricular system at a lower rate in Alzheimer’s disease compared to healthy controls. The results from the current work also indicate that the reduction of tracer entering and leaving the lateral ventricles in patient groups is independent of A*β* deposition, as indicated by the lateral ventricle PET results from the multiple sclerosis dataset that is not expected to have significant A*β* accumulation in tissue. This is further supported by our observation of no significant difference in ^11^C-PiB signal in gray matter of multiple sclerosis patients compared to controls, in contrast to the higher ^11^C-PiB signal observed in Alzheimer’s disease gray matter. All together, these results suggest that CSF-mediated tissue clearance is reduced in Alzheimer’s disease and multiple sclerosis compared to healthy controls.

Increasing age is considered one of the major risk factors in the development of Alzheimer’s disease (*44*) and A*β* deposition has been observed to be associated with increasing age in cognitively normal individuals without Alzheimer’s disease (*45*). Our observation of an inverse relationship between gray matter and lateral ventricle ^11^C-PiB signal in our mild cognitive impairment and matched control groups could indicate that tissue CSF clearance alterations are associated with, and may contribute to, A*β* deposition prior to Alzheimer’s disease onset. Under the pathogenic conditions of Alzheimer’s disease, there are likely additional factors related to the disease that contribute to further A*β* deposition (*46*), which weakens the linear relationship between CSF clearance and tissue A*β* deposition. This could explain why we did not observe the inverse relationship between CSF clearance and A*β* deposition measures in the Alzheimer’s disease group. However, due to the greater heterogeneity in tissue ^11^C-PiB measures in the Alzheimer’s disease and mild cognitive impairment groups, we may be lacking the statistical power necessary to measure the correlations in these groups. Additional mechanistic analysis with a larger cohort would be helpful in drawing further conclusions about these results. We did not observe an inverse relationship between gray matter and lateral ventricle ^11^C-PiB in multiple sclerosis or multiple sclerosis-matched controls likely because of the younger age of the cohort and/or different mechanism of the disease that does not typically involve A*β* accumulation. The positive correlation between lesion load and ^11^C-PiB signal in the multiple sclerosis group may indicate that CSF clearance alterations play a role in ongoing disease activity in multiple sclerosis.

Previous literature has reported observations of decreased A*β* clearance from the brain in Alzheimer’s disease (*47*) but it has been unclear whether this decrease is due to reduced membrane-transport from tissue to the CSF or from reduced clearance of the CSF through the CSF system. By using compartmental modelling analysis, we are able to make further inferences regarding where the changes in A*β* clearance are occurring and which tissue-types and systems may be responsible. From our results, we see that the change in A*β* clearance in Alzheimer’s disease is most likely due to both reduced clearance of A*β* from the tissue to the CSF and also due to reduced clearance of CSF through the CSF system. It is still unclear whether alterations to one of these processes may have preceded alterations to the other. Additional compartmental modelling analyses using data from other patient groups (including, but not limited to, multiple sclerosis, clinically isolated syndrome, mild cognitive impairment, and subjects with first signs of A*β* accumulation) would be useful for drawing further conclusions about our results.

The explanation as to why CSF-mediated clearance is reduced in neurological diseases such as Alzheimer’s disease and multiple sclerosis is not entirely clear. Previous research has shown that sleep is important for glymphatic system function and consequently the clearance of waste from the brain (*48*). Sleep disorders are common in both Alzheimer’s disease (*49*) and multiple sclerosis (*50*) and may play a part in the onset of disease and likely contribute to the ongoing disease processes. The symptoms of disease can also contribute to poor sleep, which could further aggravate the disease. Additionally, synaptic activity has been linked to increased levels of A*β* in ISF (*51*) and voluntary exercise has been shown to increase clearance of A*β* by the glymphatic system in aged mice (*52*). These findings suggest that physical activity also plays a role in brain clearance. Further, physical activity has been shown to be effective in preventing the onset and improving outcomes of Alzheimer’s disease (*53*) and may also contribute to improved sleep quality in health (*54*) and neurological disease (*55,56*). This provides an additional link to how sleep contributes to improved brain clearance. Previous work has shown reduced sleep quality before the onset of cognitive decline in Alzheimer’s disease (*57*). However, additional research is still required to investigate whether inactivity and/or sleep disturbances precede the onset of disease, which may help further explain the initial pathophysiology that leads to disease onset.

Additional explanations for alterations in CSF-mediated clearance in aged and Alzheimer’s disease brain could be attributed to changes in the barriers that exist between brain tissue, blood, and CSF. Age is associated with cellular atrophy and decreased cell-height of the choroid plexuses, which is exacerbated in Alzheimer’s disease (*58*). The production of CSF (*59*) and removal of solutes from the CSF (*60*) by the choroid plexuses are active processes and the cells become less energy efficient with age (*61*). Age-related changes to the cells of the choroid plexuses contribute to decreased clearance of solutes to the blood across the choroid plexuses as well as decreased CSF production, resulting in slower CSF turnover and reduced CSF-mediated brain clearance. The ependymal cells that line the ventricular system also flatten with age and exhibit greater dispersion in cilia expression (*62*). Cilia contribute to flow of CSF and communication across the ependymal layer (*63*) and the age-related changes to the ependymal layer likely also contribute to decreased CSF-mediated brain clearance.

Changes to the barriers of the ventricular system have also been observed in multiple sclerosis. A previous MRI study revealed irregularities in the ependymal layer in early multiple sclerosis that have been attributed to ependymal perivenular inflammation (*64*). Inflammation of the choroid plexuses is also common in the different forms of multiple sclerosis and at various stages of disease progression (*65,66*). The choroid plexuses could be responsible for the initial antigen presentation associated with disease onset. Under healthy conditions, immune surveillance of the brain is in part accomplished by lymphocyte entry to the CSF through the choroid plexuses (*67*). Initial entry of reactive lymphocytes through the choroid plexuses was observed before lymphocyte infiltration into brain tissue across the blood brain barrier in a mouse model of multiple sclerosis (*68*), which provides further support for the involvement of the choroid plexuses in multiple sclerosis disease onset. The alterations to the ventricular barriers in multiple sclerosis could be both a cause and a consequence of impaired CSF-mediated clearance mechanisms and further work is required to better understand the relationship and chronology of these changes.

Additional work will be done to investigate whether our ^11^C-PiB PET results are specific to Alzheimer’s disease and multiple sclerosis or whether CSF clearance is also altered in other neurological diseases. Further, we will look into use of other dynamic PET tracers and integration of quantitative MRI measures for assessing CSF clearance. It will also be helpful to explore how manipulation of known glymphatic system modulators affects the dynamic PET measures, which will allow for more informed interpretations of the results given from these PET measures in the future.

### Limitations

There are some limitations to this work. Although we still observed statistically significant differences between groups when including ventricle size as a covariate in the AD/MCI and multiple sclerosis datasets, this may not be an appropriate way to correct for ventricle size. Larger ventricle size is an inherent feature of Alzheimer’s disease and we do observe significant group differences in ventricle size across the Alzheimer’s disease, mild cognitive impairment, and healthy control groups in the current analysis (p=0.041), most notably between Alzheimer’s disease and healthy control subjects (p=0.001). Simply adding ventricle size as a covariate may, therefore, be statistically inappropriate for use in our model because we are likely removing variance between groups that exists due to disease. Additional work should be performed, possibly including revised inclusion criteria and group matching, to investigate more appropriate ways to accommodate for differences in ventricle size in the future. We are also restricted in our compartmental modelling analysis due to absent structural image data in the form of MRI or CT for the selected Alzheimer’s disease dataset. We produced all lateral ventricle ROIs for the compartmental modelling Alzheimer’s disease dataset using a registered MNI atlas as a template for each subject. Registered MNI atlases are not as reliable for defining brain structures as MRI or CT images and therefore our lateral ventricle ROIs for compartmental analysis are prone to error. Our selection of ^11^C-PiB PET datasets was restricted to those that included an arterial input function that is required for compartmental modelling analysis. However, the invasive nature of continuous arterial blood sampling discourages use of arterial input functions in human studies. Image-derived arterial input functions have been developed (*69*) that seemingly eliminate the need for invasive blood sampling. Unfortunately, these image-derived arterial input functions cannot be reliably used in compartmental modelling analysis. Ideally, future PET studies will be designed to include arterial blood sampling during PET data collection for specific use in compartmental modelling analysis. Further, the same compartmental modelling analysis should be performed on a multiple sclerosis dataset with PET data collected with an arterial input function to determine whether the mechanism behind the change in CSF dynamics is shared between Alzheimer’s disease and multiple sclerosis. Our compartmental model is also limited in that it uses TAC data from all gray matter for the tissue pool that exchanges with the lateral ventricles. We only expect direct exchange between the deep gray matter structures and the lateral ventricular CSF because of their proximity to each other. However, these regions of direct exchange are difficult to define and signal from whole gray matter is expected to behave similarly to the signal in the deep gray matter structures alone. Therefore, signal from whole gray matter was used as a representative measure for the gray matter that exchanges with the lateral ventricle CSF. We also tested a model using signal from cerebellar gray matter as a region that is not expected to have specific binding in Alzheimer’s disease (Supplementary Fig. 1) to confirm that the differences in rate constants between groups is not driven by differences in gray matter signal due to specific binding to Aβ. A model that entirely excluded input from the gray matter tissue to the lateral ventricles (Supplementary Fig. 2) resulted in poorer fits of the lateral ventricle TAC data, indicating that this simplified model is inferior in its ability to represent the system. Finally, it is unclear what the specific binding in the lateral ventricles represents in our model. This bound pool may be accounted for by tracer binding to the ventricle walls and/or choroid plexus or, less likely, to compounds within the CSF. We found that a model that excluded the ventricular bound pool (Supplementary Fig. 3) did not reliably fit the lateral ventricle TAC data, indicating that this simplified model does not accurately represent the system. Inclusion of a ventricular bound pool in our final model is also consistent with a recently presented model of the CSF clearance system (*42*).

### Conclusion

The results from this work confirm that dynamic ^11^C-PiB PET can be used to assess CSF dynamics in health and neurological disease. Although it is not yet confirmed whether we are also measuring glymphatic activity, the close connection between the CSF and glymphatic systems, as well as our results showing dynamic PET differences in patient groups with known glymphatic dysfunction, suggest that this method might be used to assess glymphatic function after further validation. Our results provide further support for a promising method for assessing brain clearance in neurological disease. We hope that this and future work in this area will improve the understanding of the pathogenesis of neurological diseases such as Alzheimer’s disease and multiple sclerosis.

## Supporting information

## Acknowledgements

We would like to acknowledge Hammersmith Imanet for provision of radiotracers and scanning facilities used for acquiring the Alzheimer’s disease and healthy control data for the AD/MCI dataset. Amyloid tracer used for the AD/MCI dataset was made available by GE Healthcare. Additionally, we would like to thank Hope McDevitt, Stella Ahier, Andreanna Williams, James Anscombe, and Andrew Blyth for assisting with scanning of the AD/MCI cohort. We thank ASREP for supporting B.B., the Centre d’Investigation Clinique team from ICM and Jean-Christophe Corvol for protocol organization as well as C. Baron, P. Bodilis (CEA), C. Dongmo, and G. Edouart for assistance with the multiple sclerosis dataset, and Ramin Parsey from the Columbia PET centre for use of the PET data used in our compartmental modelling analysis. The original study that collected the Alzheimer’s disease data used in our compartmental modelling analysis received research support from GlaxoSmithKline. We would also like to graciously acknowledge all study participants that took part in the studies included in this work.

## Abbreviations

A*β*: = amyloid-beta
AUC: = area-under-curve
AQP4: = aquaporin-4
EDSS: = Expanded Disability Status Scale
ISF: = interstitial fluid
MSSS: = Multiple Sclerosis Severity Score
ROI: = region of interest
SUVR: = standard uptake value ratio
TAC: = time activity curve

