## Supplementary material for "Dynamic 11C-PiB PET shows cerebrospinal fluid flow alterations in Alzheimer’s disease and multiple sclerosis"

Julia J. Schubert et al.

### Supplementary material

*Testing of compartmental models*

Model 1: Cerebellar gray matter as a region not expected to have specific tracer binding used to represent tissue input to lateral ventricles to confirm that differences in rate constants is not due to amyloid-beta accumulation in tissue.

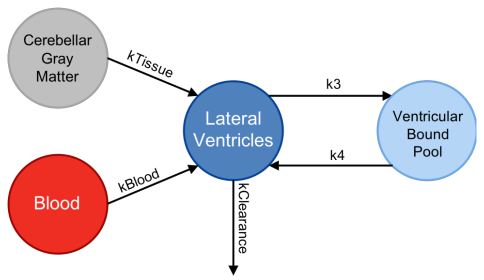


**Supplemental Figure 1. Compartmental model using only cerebellar gray matter as kTissue input to lateral ventricles.** *Kclearance* includes total clearance from lateral ventricles to blood, tissue, and rest of ventricular system.

Model 2: Simplified model that excludes input from tissue to the lateral ventricles.

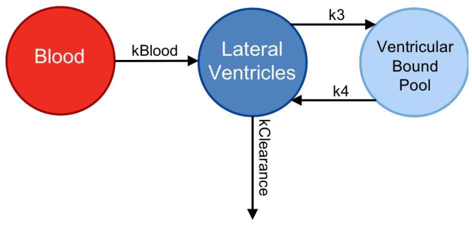


**Supplemental Figure 2. Simplified model excluding gray matter tissue input to lateral ventricles.** *Kclearance* includes total clearance from lateral ventricles to blood, tissue, and rest of ventricular system.

Model 3: Simplified model that excludes ventricular bound pool.

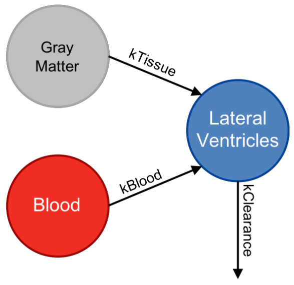


**Supplemental Figure 3. Simplified model excluding ventricular bound pool.** *Kclearance* includes total clearance from lateral ventricles to blood, tissue, and rest of ventricular system.

Model 4: Final model used in analysis that uses whole gray matter as tissue input to lateral ventricles and includes ventricular bound pool.

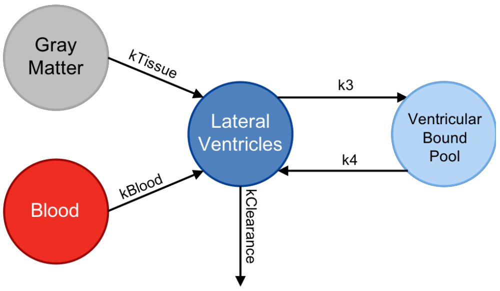


**Supplemental Figure 4. Final model used in compartmental modelling analysis.** *Kclearance* includes total clearance from lateral ventricles to blood, tissue, and rest of ventricular
